# HGC: fast hierarchical clustering for large-scale single-cell data

**DOI:** 10.1101/2021.02.07.430106

**Authors:** Ziheng Zou, Kui Hua, Xuegong Zhang

**Affiliations:** MOE Key Laboratory of Bioinformatics, Bioinformatics Division, BNRIST and Department of Automation, Tsinghua University, Beijing 100084, China; Center for Synthetic and Systems Biology and School of Medicine, Tsinghua University, Beijing 100084, China

## Abstract

Clustering is a key step in revealing heterogeneities in single-cell data. Cell heterogeneity can be explored at different resolutions and the resulted varying cell states are inherently nested. However, most existing single-cell clustering methods output a fixed number of clusters without the hierarchical information. Classical hierarchical clustering provides dendrogram of cells, but cannot scale to large datasets due to the high computational complexity. We present HGC, a fast **<u>H</u>**ierarchical **<u>G</u>**raph-based **<u>C</u>**lustering method to address both problems. It combines the advantages of graph-based clustering and hierarchical clustering. On the shared nearest neighbor graph of cells, HGC constructs the hierarchical tree with linear time complexity. Experiments showed that HGC enables multiresolution exploration of the biological hierarchy underlying the data, achieves state-of-the-art accuracy on benchmark data, and can scale to large datasets. HGC is freely available for academic use at https://www.github.com/XuegongLab/HGC.

## Introduction

Recent developments of single-cell RNA sequencing (scRNA-seq) technologies and bioinformatic tools have accelerated our understanding of cell heterogeneity [1,2]. A fundamental task in scRNA-seq data analysis is to cluster cells into different groups as candidate cell types or cell states. This task is difficult due to the challenging characteristics of scRNA-seq data and the complex nature of cell heterogeneity underlying the data [3]. One of the common structures of cell heterogeneity is the hierarchical structure, which can be explored at different resolutions and results in varying and nested cell types or cell states. Current practice for studying such multi-level heterogeneity is to first produce a fixed number of clusters and then adjust the clustering resolutions in an *ad hoc* manner [3,4]. This workflow loses the underlying hierarchical information and requires multi-rounds of re-clustering to find a suitable resolution. As an alternative, hierarchical clustering enables direct multi-resolution exploration of the hierarchical cell heterogeneity. However, classic hierarchical clustering algorithms are only suitable for small datasets due to the high computational complexity.

We propose a fast Hierarchical Graph Clustering method HGC for large-scale single-cell data. The key idea of HGC is to construct a dendrogram of cells on their shared nearest neighbor (SNN) graph. This combines the advantages of graph-based clustering methods and hierarchical clustering. We applied HGC on both synthetic and real scRNA-seq datasets. Results showed that HGC can recover the biological hierarchy underlying the data, can achieve high clustering accuracy at fixed resolution, and can scale well to large datasets.

## Methods

The workflow of HGC contains two major steps: graph construction and dendrogram construction. For the graph construction step, HGC adopts the standard procedure of building the SNN graph, which is to first conduct Principal Component Analysis (PCA) on the expression data and build the k nearest neighbor (KNN) graph and the SNN graph in the PC space (**Fig. 1**) [5]. The construction of dendrogram on the graph is a recursive procedure of finding the nearest neighbor pair and updating graph by merging node pairs (**Fig. 1**).

**Fig. 1.**
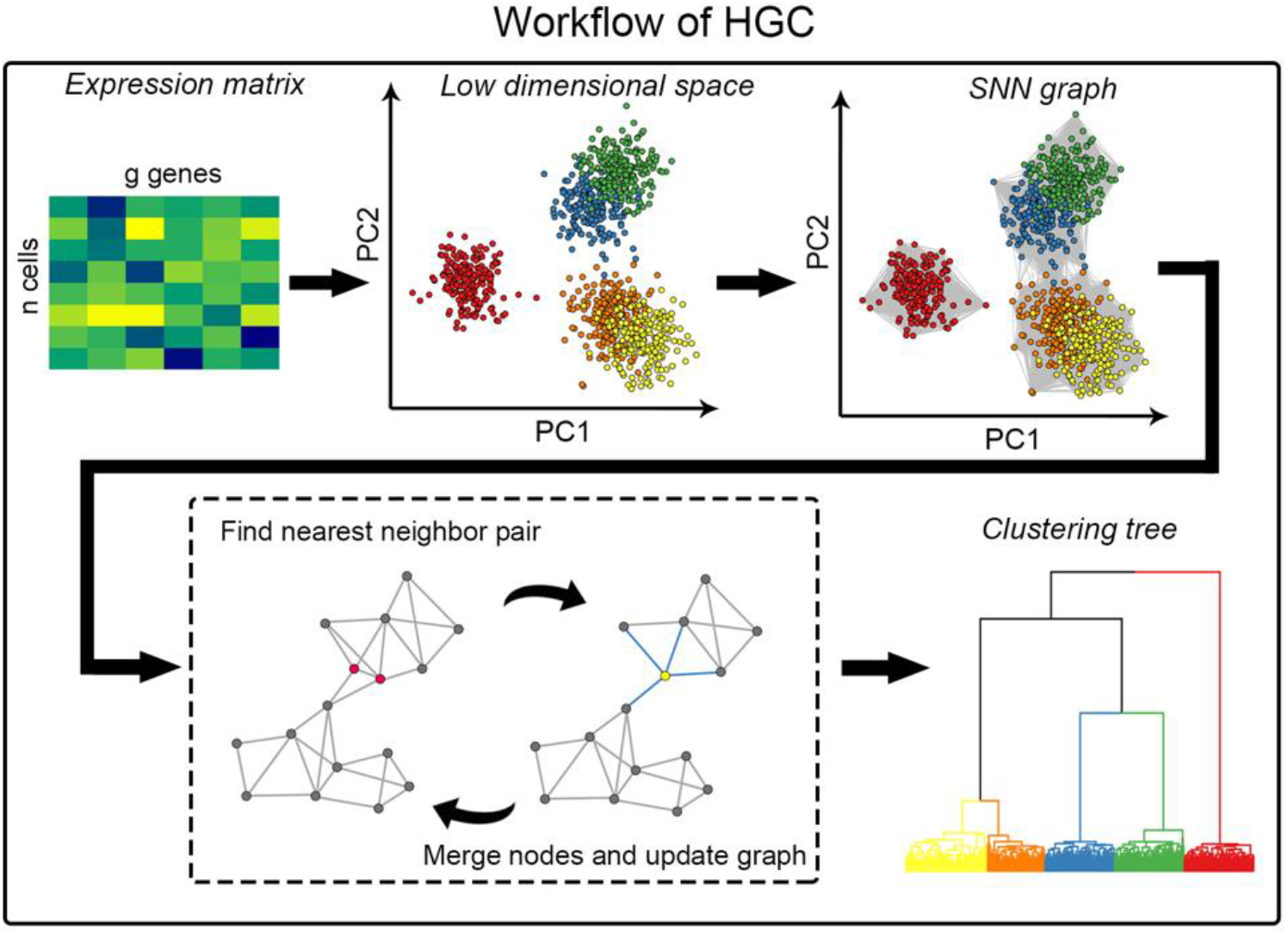
The workflow of HGC. HGC contains two main steps, the construction of SNN graph in the PC space, and a recursive procedure of finding the nearest-neighbor node pairs and updating the graph by merging the node pairs. HGC outputs a dendrogram like classical hierarchical clustering.

The key to find the nearest neighbor pair on a graph is the distance measure. HGC utilizes the node pair sampling distance introduced in [6]. For a weighted, undirected graph *G* = (*V, E*), let *A* be the weighted adjacent matrix. If we sample node pairs or edges at random in proportion to their weights, the probability that node pair or edge (*i, j*) being sampled is:

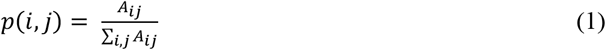

Similarly, sampling nodes in proportion to their weighted degrees results in the probability of node *i* being sampled:

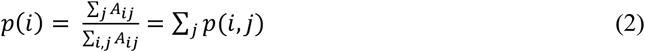

The node pair sampling distance is then defined as the ratio between individual sampling probability and the pair sampling probability:

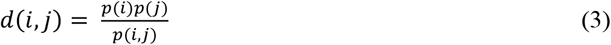

After finding two nearest neighbor nodes, the graph is updated by merging them into a new node (**Fig. 1**). Such new nodes in the updated graph are communities of the original nodes. The weighted degree of the new nodes is the sum of the weighted degrees of all original nodes in the community. Weights of edges between new nodes are the sum of the weights of the edges between original nodes in the corresponding two communities.

The node pair sampling distance has been proven to have the property of reducibility [6], which enables the whole hierarchical clustering procedure to be accelerated using the nearest neighbor chain algorithm. This algorithm searches the mutual nearest neighbors instead of the closest pairs, resulting in the same dendrogram as the standard searching procedure in a much shorter time.

The graph construction step in HGC is flexible. Besides the SNN graph used in the default workflow, one can use other types of graphs, such as the KNN graph, or graphs built from other single-cell tools. The preprocessing in the graph construction can also be adjusted. In the benchmark experiments, we have included the results of using GLMPCA as the dimension reduction methods instead of PCA.

HGC has been implemented in R, with the key function written in Rcpp. We provide Seurat-style function to guarantee seamlessly usage in this popular pipeline. It also includes tools to assist downstream analysis, such as the dynamicTreeCut package for cutting the dendrogram into specific clusters, and plotting functions for visualizing the hierarchical clustering results.

## Results

### HGC reveals hierarchical structure of cell heterogeneity

We applied HGC on two datasets with known hierarchical structures: the Pollen dataset from [7] and the PBMC dataset from [8]. As the baseline, we experimented classical hierarchical clustering methods on these two datasets. The pairwise distance required in classical hierarchical clustering can be calculated in different feature spaces. We considered three feature spaces: the gene expression space and the feature spaces given by applying PCA or GLMPCA [9] on the gene expression data. The three corresponding methods are referred to as HC, PCA+HC and GLMPCA+HC, respectively. They are called HC-based methods when referred to together.

### Experiments on the Pollen dataset

Cells in the Pollen dataset can be classified at two levels: the tissue level and the cell line level [7]. At the tissue level, they can be divided into 4 groups according their tissue source, including blood cells, dermal cells, human-induced stem cells, and nerve cells. Each of these groups can be further divided into subgroups. Blood cells include K562, HL60 and CRL-2339 cell lines. K562 cell line and HL60 cell line are from leukemia patients. CRL-2339 cell line is B lymphoblasts. Dermal cells include Kera cells, BJ cells and CRL-2338 cells. Kera cells are a type of skin cell line. The BJ cell line is a human fibroblast, and the CRL-2338 cell line is an epithelial cell derived from ductal carcinoma. Human-induced pluripotent cell (hiPSC) is considered a single tissue, which is derived from the BJ cell line. Neurons include Neural Progenitor Cells (NPC), GW16 cells, GW21 cells and GW21+3 cells. NPC is derived from human induced pluripotent stem cells. GW cells are cells in the human genital area from different embryonic developmental stages.

For the Pollen dataset, HGC gave a reasonable clustering result. As shown in **Fig. 2a**, HGC detected five main clusters. In the SNN graph, these five classes were not connected to each other, so HGC cannot further merge them. This clustering result is slightly different from the classification at the tissue level (ARI=0.61). Neurons and hiPSC were grouped into one cluster, which can be explained by their differentiation relationship. Dermal cells were assigned to BJ cells and non-BJ cells and blood cells were clustered as the K562 and non-K562 group. When a larger k is used to split the hierarchical clustering tree, these main tissue sources will be further divided into smaller clusters. When k = 11, almost all 11 cell lines were identified by HGC (ARI = 0.94, **Fig. 2c**). The hierarchical relationship of these cell lines according to the dendrogram by HGC is shown in **Fig. 2d,** which agrees well with the prior knowledge.

**Fig. 2.**
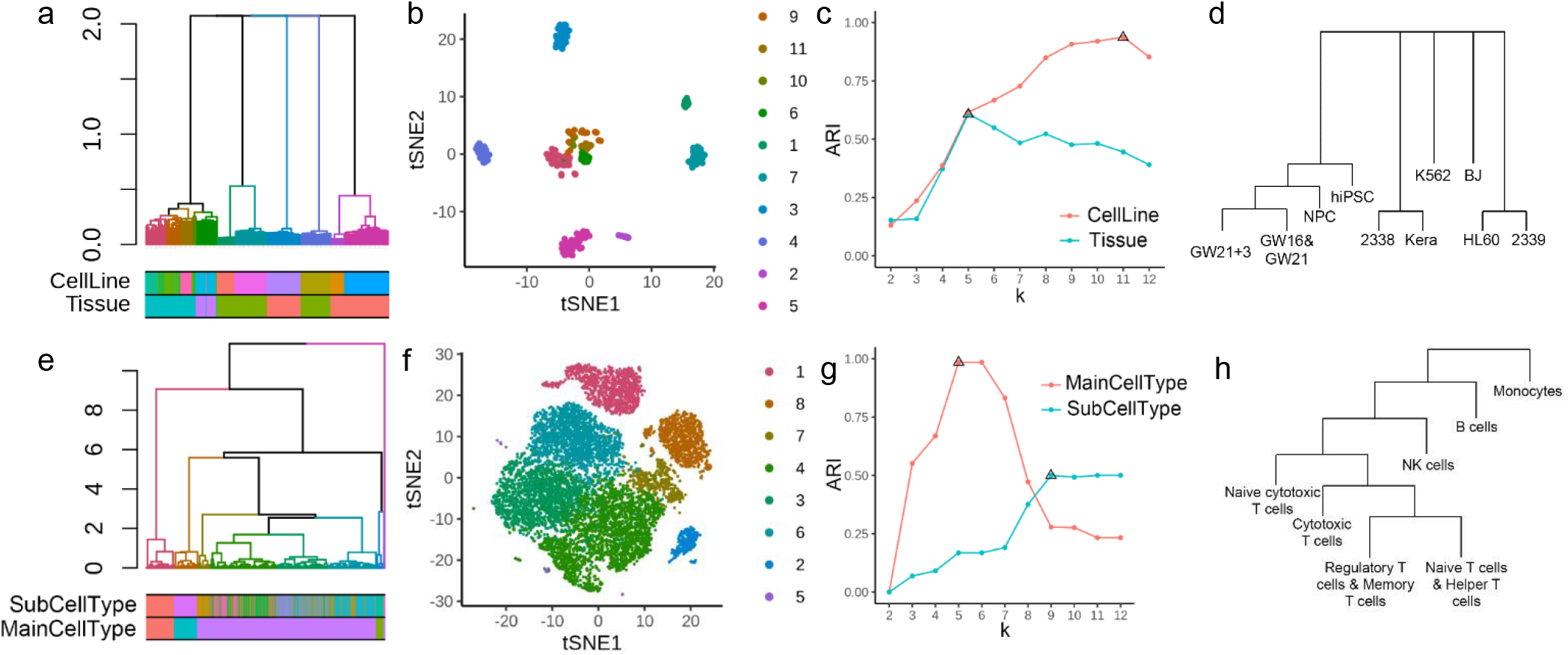
The clustering trees by HGC capture the multi-layer structure in datasets. (a) The dendrogram given by HGC for Pollen dataset. The color bars below the dendrogram are the two-level labels. (b) The tSNE plot showing the clustering result when cutting the tree into 11 clusters. (c) The ARIs of the clustering results compared with two labels. The x-axis is the number of clusters and the triangles represent the maximal ARIs for the two labels. (d) The inferred hierarchical relationship of different cell lines the Pollen dataset given by HGC. (e-h) The corresponding results for PBMC dataset.

As the baseline, we ran the three HC-based methods on the Pollen dataset (**Fig. S1**). We cut the dendrogram into different numbers of clusters and calculated the ARI between the clusters and the two labels (**Fig. S1**). For both labels, the three HC-based did not give clustering results that agree well with the labels. Visualization of the dendrograms showed the reasons of the bad performance. The hierarchical trees given by the HC-based methods tended to form many small branches, leading to poor clustering results when cutting the tree into specific clusters. Taking the result of GLMPCA+HC as an example, it can be seen from the annotation result of the color barcode that the clustering result captures certain clustering information. However, dendrogram had many small branches. When cutting the tree into, say 11 clusters, these small branches formed individual clusters, and 80% of the cells are classified into the same cluster, resulting in an ARI of only 0.02.

Besides hierarchical clustering, current practice to reveal the multi-layer heterogeneity is to conduct multi-rounds of clustering with different resolutions. We experimented this strategy here using Seurat. In Pollen dataset, a resolution of 0.8 in Seurat gave similar results to that of HGC when k = 5. When the resolution ranged from 1 to 3, Seurat gradually gave finer clustering results. At a resolution of 3, Seurat recovered almost all 11 cell lines. When the resolution increases, Seurat tends to cut the existing clusters into finer ones, reflecting certain level of cell hierarchy. However, such partial hierarchical information is empirical and only available with the assistance of other visualization results such as tSNE or UMAP.

### Experiments on the PBMC dataset

The PBMC dataset contains 9 different immune cell types obtained through FACS, including B cells, monocytes, NK cells and 6 subtypes of T cell (**Table S1**). The differentiation relationship of immune cells is relatively clear. For the cell types included in the PBMC dataset, monocytes are derived from myeloid progenitor cells, while T cells, B cells and NK cells are derived from lymphoid progenitors. T cells can be further divided into two major subtypes: CD4^+^ T cells and CD8^+^ T cells.

HGC obtained a biologically meaningful multi-level structure in the PBMC dataset. When k=5, HGC captured four main cell types in the data (ARI=0.98). Among them, a small part of monocytes was grouped into a single class as a new branch, which is consistent with the results of tSNE plot (**Fig. 2e,f**). When k increased from 5 to 9, new clusters were generally generated inside T cells. HGC discovered four main branches of T cells: the naive cytotoxic T cells branch, cytotoxic T cells branch, a branch of the mixture of regulatory T cells and memory T cells, and a branch of the mixture of naive T cells and helper T cells. From the dendrogram given by HGC, we can roughly infer the hierarchical relationship of each cell type. The cell types deduced from roots to leaves are monocytes, B cells, NK cells, CD8^+^ T cells, and finally CD4^+^ T cells (**Fig. 2h**), which is consistent with biological knowledge. The four subtypes of CD4^+^ T cells are more difficult to be classified based on transcriptional data, which is also consistent with the current understanding of T cell subtypes [10].

As comparisons, we ran the HC-based methods on the PBMC dataset. The classical hierarchical clustering algorithm has a relatively high computational complexity. Since the purpose here is to examine the ability to capture the hierarchical structure rather than the efficiency, we used the “geometric sketching” algorithm to down-sample the data (**Table S1**) [11]. The results of the three HC-based methods are shown in **Fig. S2**. HC separated part of the NK cells, B cells, and monocytes, but clustered the remaining B cells, NK cells, monocytes and T cells together (**Fig. S2a**). In the task of distinguishing the main cell types, PCA+HC and GLMPCA+HC both performed better than HC. But like what we have discussed in the Pollen dataset, the HC-based methods generated many small branches, and most of the cells were clustered into one large branch. This made it hard to determine clusters based on the hierarchical clustering results.

We also used Seurat to obtain a series of clustering results with different resolutions. At resolution of 0.01, Seurat identified four main clusters, which correspond to T cells, NK cells, B cells and monocytes. With the increase of resolution, Seurat first separated cytotoxic T cells and naive cytotoxic T cells from the main cluster of T cells. Then it identified the cluster of the initial T cells and regulatory T cells. Finally Seurat split the cluster of B cells and monocytes to find finer subgroups when resolution is 1.0.

### Comparison with existing methods on benchmark datasets

To further benchmark HGC’s performance on revealing cell heterogeneity at fixed level, we collected six scRNA-seq datasets with known labels and compared 15 existing clustering methods with adjusted rand index (ARI) and normalized mutual information (NMI) between clustering results and known labels. Detailed information about the evaluated methods, the two evaluation indexes and the benchmark datasets are introduced as follows.

### Existing methods compared

We collected the state-of-the-art clustering methods to compare with HGC, including Seurat, SC3, monocle3, TooManyCells, CIDR, SIMLR, RaceID3, CountClust and densitycut [3,5,12–18]. We also included the three HC-based methods, HC, PCA+HC and GLMPCA+HC, as the baseline. Seurat is one of the most popular scRNA-seq data processing pipelines. The default clustering method in Seurat is to conduct the Louvain algorithm on the SNN graph built in the PC space given by PCA [5]. We also evaluated building the SNN graph in the PC space given by GLMPCA before Louvian clustering, which we referred to as Seurat_GLMPCA. Similarly, HGC using PCA and GLMPCA as the dimension reduction method are noted as PCA+HGC and GLMPCA+HGC, separately. SC3 is a consensus clustering method, which first applies kmeans with different distance metrices and preprocessing methods, and then achieves a consensus clustering result using classical hierarchical clustering [3]. For large datasets, SC3 first achieves clustering results on a small subset of the datasets, and using the clustering results as labels to train an SVM to classify the remaining cells. Monocle3 is a state-of-the-art trajectory inference pipeline, which includes the Louvain algorithm the clustering algorithm and obtain the clusters on the cell graph built in the UMAP space [12]. TooManyCells is a divisive hierarchical clustering workflow to catch and visualize the cell clades. It partitions the cells into two groups with the spectral clustering algorithm in an iterative manner. TooManyCells does not give the full hierarchy and the stopping criterion is determined based on the Newman–Girvan modularity [13]. CIDR first conducts a dropout-aware dimension reduction on the expression data and then applies the classical ward hierarchical algorithm in the low-dimension space [14]. SIMLR first learns a proper cell-cell distance metric using multiple kernels, and then conducts downstream visualization and clustering based on the metric [15]. RaceID is a method specially designed for detecting rare cell types in scRNA-seq data. It treats this task as an outlier detection problem and solves it using k-means [16]. CountClust adopts the topic model in natural language processing as the clustering method where topics are treated as clusters [17]. Densitycut is a density-based clustering algorithm. It estimates the local densities for each data point and detects the density peaks in the dataset as clusters [18].

For a fair comparison, all parameters in these methods were set as the default values except the parameter related to cluster number. For methods that need to specify the cluster number, the known cluster numbers in benchmark datasets were used. Methods that do not require a specific cluster number, such as densitycut, Seurat and monocle3, were run with default parameters. For SC3, we used it on all cells instead of training an SVM model. For TooManyCells, it does not need the cluster number and can decide when to stop splitting the clades. However, the divisive nature of TooManyCells makes it hard to set as specific cluster number. We chose the layer which gave the closest cluster numbers to the known ones.

### Evaluation indexes

To evaluate the performance of clustering methods, we adopted the ARI and NMI to measure the agreement between the clustering result and the known label.

Rand Index (RI) is a measure of the similarity between two clustering results on one dataset, and ARI is the corrected-for-chance version of RI. RI is calculated based on a pairwise comparison between two clustering results. The calculation of RI is to first enumerate all pairs of points in the datasets and record if two clusterings give the same results on them. RI is then calculated as the fraction of all pairs that two clustering results agree with each other:

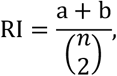

where n is the number of points (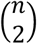 is therefore all possible pairs of points), a is the number of pairs assigned to the same cluster in both clusterings, and b is the number of pairs assigned to different clusters in both clusterings.

ARI is the corrected-for-chance version of RI. Such a correction for chance establishes a baseline by using the expected similarity of all pair-wise comparisons between clusterings specified by a random model. Briefly, the definition of ARI is

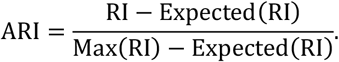

There are different selections for the random model. We adopted the most widely used permutation model in our experiments.

NMI measures the similarity using information theory. The mutual information describes the dependence between two variables, which here are two clustering results. The mutual information could be defined with the entropy. Denote the two clustering results as Y and C, the entropy of clustering Y is defined as

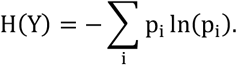

where the p_i_ is the fraction of points assigned to cluster i in clustering Y. The mutual information is defined as

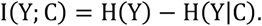

H(Y|C) is defined with the help of conditional distributions. And the NMI is

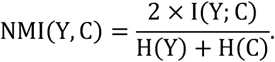

Mutual information could take a wide range, while the NMI’s value is limited in [-1,1]. Larger NMI value indicates high similarity.

### Benchmark datasets and results

We collected six benchmark datasets, including two synthetic datasets and four real datasets. Detailed information and results on them are introduced as follows. Overall, HGC, Seurat and SC3 achieved comparable performance and outperformed other methods (**Fig. 3**). Other methods like monocle3 and RaceID3 also produced good performance on most of the datasets, but they were not as stable as the top three methods.

**Fig. 3.**
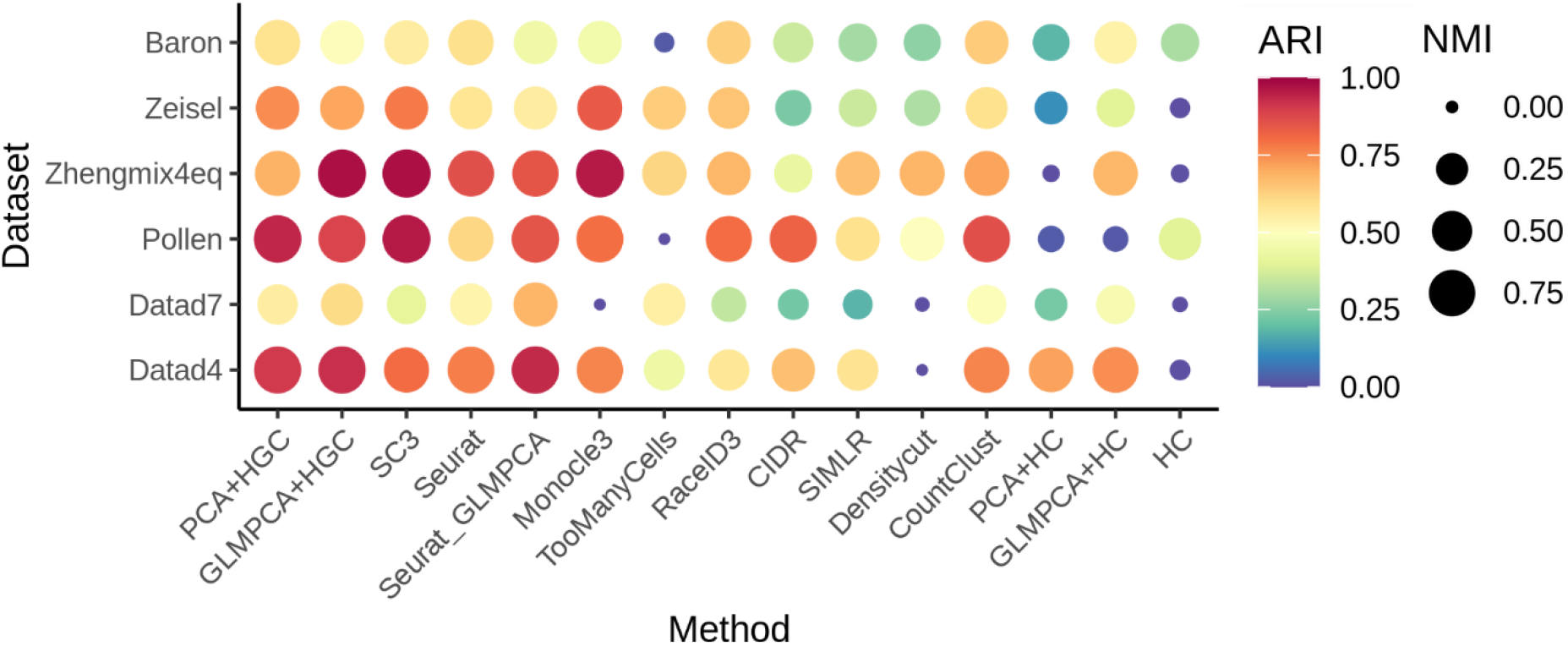
Benchmarking of 15 clustering methods on six datasets. We used ARI and NMI to measure the agreement between clusters found with the methods and the known labels. HGC, Seurat and SC3 achieved comparable clustering accuracy and significantly outperformed other methods

### Synthetic datasets

The synthetic datasets were generated using SymSim, a tool for simulating scRNA-seq data [19]. SymSim models the variation of the observed data as three parts: extrinsic variation, intrinsic variation and technical variation. Extrinsic variation refers to the variation caused by external variability factors (EVF). The model for EVFs could be discrete or continuous, with adjustable parameters for users. Intrinsic variation models the intrinsic dynamic progress of the transcription. Extrinsic variation and intrinsic variation jointly determine the true transcript counts of each cell. Technical variation adds another layer of variations in the observed transcription data caused by the differences in sample processing procedure, sequencing protocols and other technical reasons.

We experimented two synthetic datasets generated by SymSim, which we referred to as Datad4 and Datad7, respectively. They were generated with the same parameters except the different variances of EVFs. They both contain five discrete clusters and each cluster includes 200 cells.

In Datad4, PCA+HGC, GLMPCA+HGC and Seurat_GLMPCA achieved comparable accuracy which outperformed other methods. SC3, Seurat, monocle3 and CountClust also gave good clustering results. As the baseline, HC achieved an ARI of about zero, meaning that it almost totally lost the cluster information. PCA+HC and GLMPCA+HC provided better performance, suggesting the importance of dimension reduction in single-cell clustering tasks.

In Datad7, the performance of all methods dropped due to increase of noise. Seurat_GLMPCA, PCA+HGC and GLMPCA+HGC were still the top 3 methods. Seurat, CountClust and TooManyCells also gave comparable accuracy to those of the top 3 methods. With the default parameters, monocle3 assigned all cells to a single cluster which resulted in low ARI and NMI. HC-based methods again did not perform well. The rankings of the clustering methods are generally the same in the two datasets.

### Real scRNA-seq datasets

We experimented four real scRNA-seq datasets whose sizes range from hundreds to thousands of cells [7,8,20,21]. These datasets come from different tissues and various sequencing protocols (**Table S2**).

As introduced before, cells in the Pollen dataset can be classified at two levels [7]. We used the 11 cell lines as labels in the benchmarking experiments. Results showed that SC3, PCA+HGC and GLMPCA+HGC achieved the top accuracies. Seurat and monocle3 also gave good performance. It was interesting to see that HC applied directly on the expression data got higher ARI than PCA+HC and GLMPCA+HC, although their performance were the worst among all methods. TooManyCells does not accept un-integer matrix as input, therefore we did not include it when benchmarking performance in Pollen dataset.

The Zhengmix4eq dataset contains four immune cell types from [8]. The cell types are B cells, monocytes, naive cytotoxic T cells and regulatory T cells. Each cell type has the same number of cells. GLMPCA+HGC, SC3 and monocle3 well recovered the partition of the four cell types. The performance of PCA+HGC (ARI = 0.69) was not as good as GLMPCA+HGC, because when cutting the dendrogram into four clusters, it merged the two T cell types together and split the monocytes into two clusters. HC and PCA+HC got ARIs of about zeros. GLMPCA+HC got a meaningful clustering result with an ARI of 0.68, which again suggested the importance of preprocessing.

The Zeisel dataset contains cells from the mouse brain. In the original paper, the label of the cells was obtained through a biclustering clustering method called BackSPIN [20]. The nine cells types were determined by the marker genes. The clustering task for this dataset is hard because transcriptional difference among neurons and neuroglial cells is subtle. Monocle3 got the best result with ARI 0.84. PCA+HGC, GLMPCA+HGC and SC3 achieved comparable good accuracy with the ARIs ranging from 0.77 to 0.71. The three HC-based methods produced bad results, with ARIs smaller than 0.5.

The Baron dataset contains cells from human and mouse pancreas [21], and we utilized the human data in our experiments. In the original paper, cells are classified into fourteen cell types with an iterative hierarchical clustering framework. The classifications are validated by known molecular markers. ARIs and NMIs showed that monocle3, SC3 and PCA+HGC achieved the comparable accuracy that outperformed others. GLMPCA+HGC and RaceID also produced good clustering results with the ARIs around 0.7. GLMPCA+HC achieved an ARI of 0.51 and HC, PCA+HC produced bad clustering results.

### Scalability

We tested the time efficiency of HGC in the Mouse Cell Atlas (MCA) dataset which contains about 400,000 cells [22]. To reflect how the running time changes as the sample size increases, we sampled a series of datasets from MCA whose sizes range from 10,000 cells to 400,000 cells. As comparisons, we applied HC and Seurat on those datasets. For each of the three methods, we recorded the running time of preprocessing step and clustering step (**Fig. 4**). The preprocessing step refers to the calculation of pairwise distance in HC, and the construction of SNN graph in Seurat and HGC. Results showed that the construction of SNN graph is much faster than calculating the pair-wise distance, which validates the advantage of graph-based clustering in terms of efficiency. The running time of the dendrogram construction step in HGC grows almost linearly as the sample size increases, significantly outperforming both HC and Seurat clustering (**Fig. 4**).

**Fig. 4.**
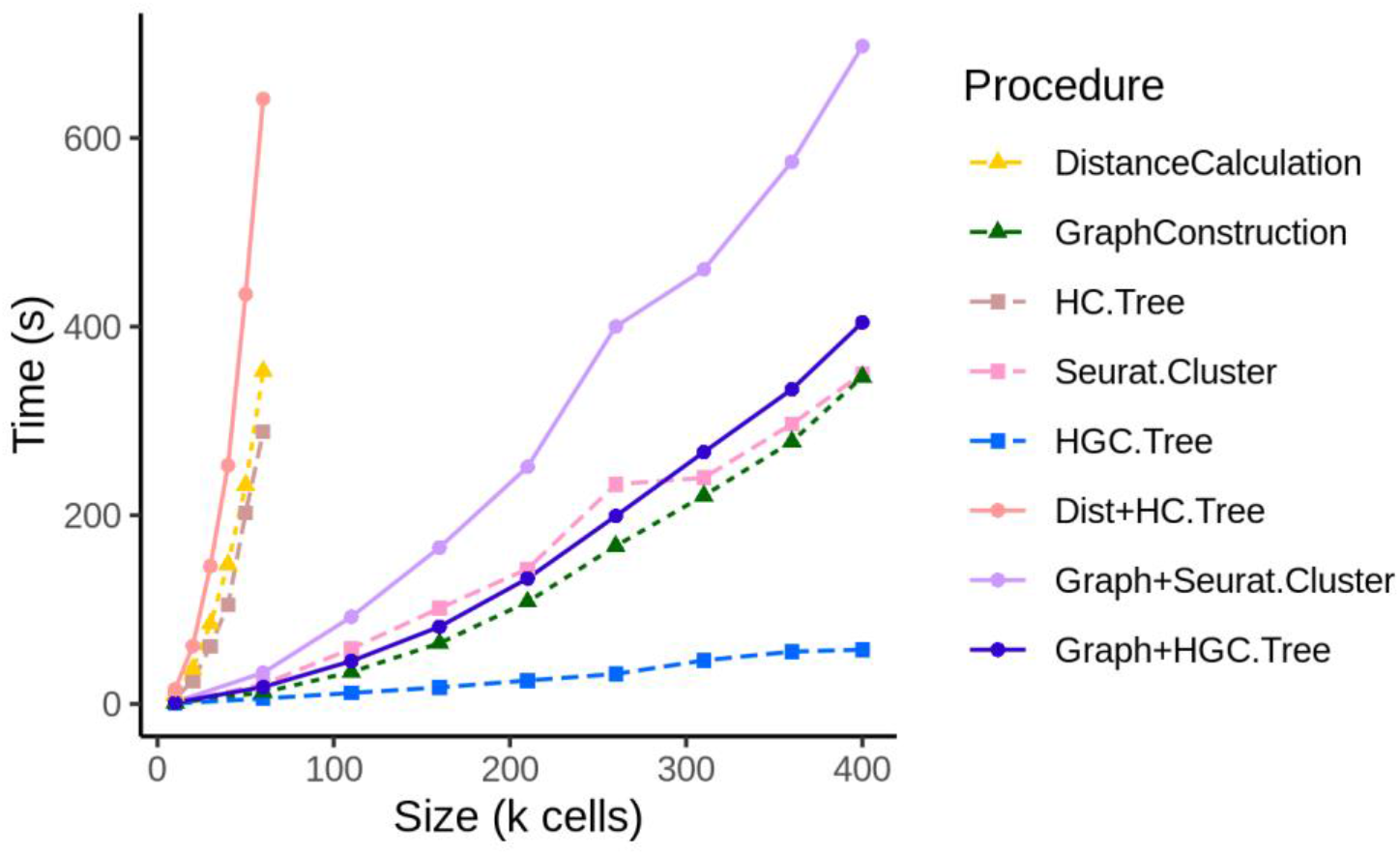
Benchmark of the running time. We tested the running time of HGC, HC and Seurat on a series of datasets with different sizes sampled from MCA. The three shapes of lines show the running time of different procedures: the triangle+dashed line represents the running time for pre-processing step, which refers to the calculation of pairwise distance in HC (DistanceCalculation) and the construction of the SNN graph in HGC and Seurat (GraphConstruction). The square+dashed line represents the running time of building the hierarchical tree (HG.Tree for classical hierarchical clustering and HGC.Tree for HGC) and running Louvain clustering in Seurat (Seurat.Cluster), and the circle+solid line represents the running of the whole clustering pipeline (Dist+HC.tree for classic hierarchical clustering, Graph+HGC.tree for HGC and Graph+Seurat.Cluster for Seurat).

### Conclusion

We developed an R package HGC for conducting fast hierarchical clustering of single-cell data dataset. HGC can recovery the hierarchical structure underlying the data that has been mostly ignored in current clustering methods. HGC achieves comparable clustering accuracy with the state-of-the-art clustering methods with a better scalability to large single-cell data. These properties make HGC a convenient tool for exploring the hierarchical heterogeneity in single-cell studies.

---

**Table S1.**
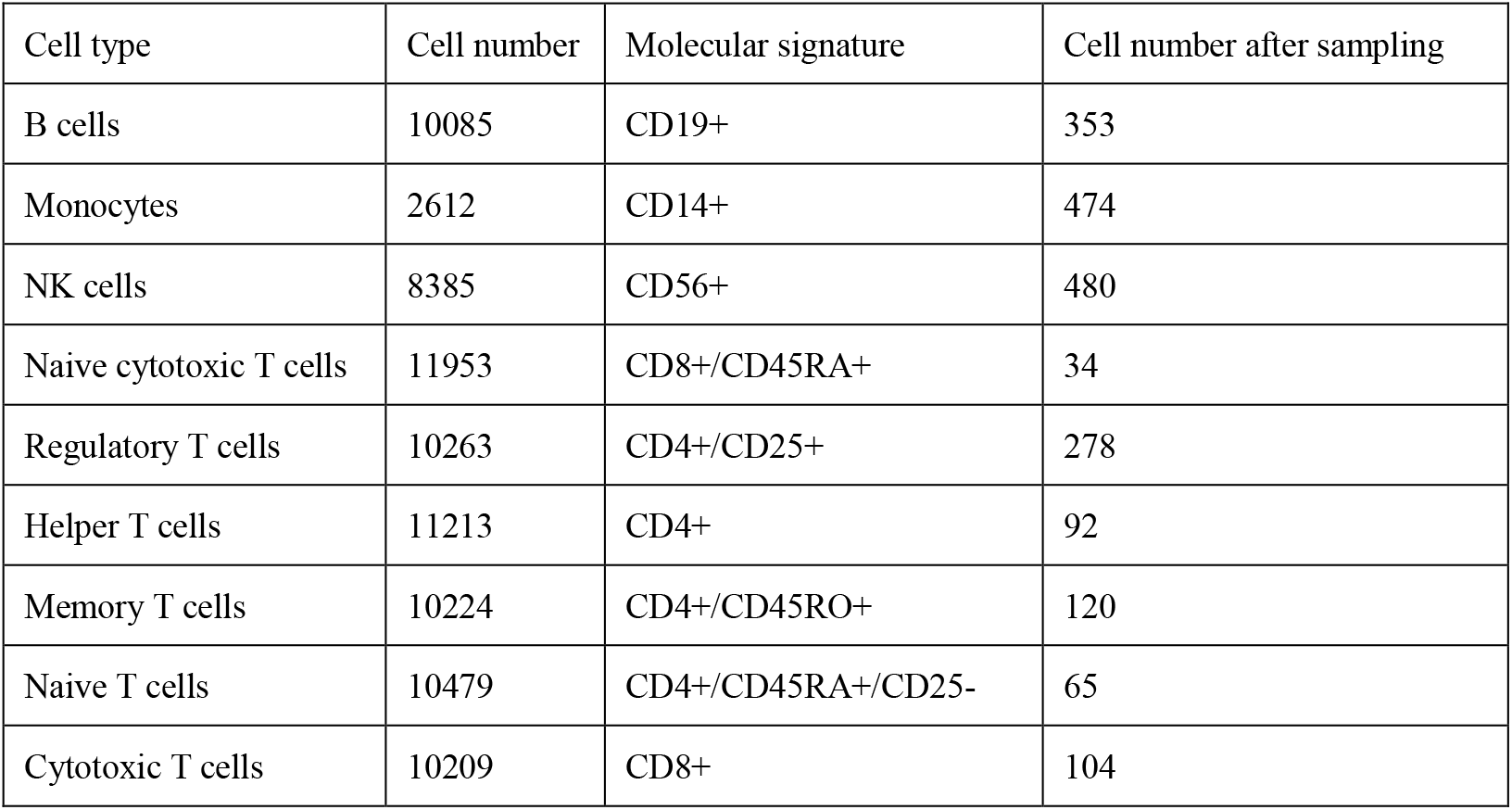
Cell types in the PBMC dataset

| Cell type | Cell number | Molecular signature | Cell number after sampling |
| --- | --- | --- | --- |
| B cells | 10085 | CD19+ | 353 |
| Monocytes | 2612 | CD14+ | 474 |
| NK cells | 8385 | CD56+ | 480 |
| Naive cytotoxic T cells | 11953 | CD8+/CD45RA+ | 34 |
| Regulatory T cells | 10263 | CD4+/CD25+ | 278 |
| Helper T cells | 11213 | CD4+ | 92 |
| Memory T cells | 10224 | CD4+/CD45RO+ | 120 |
| Naive T cells | 10479 | CD4+/CD45RA+/CD25- | 65 |
| Cytotoxic T cells | 10209 | CD8+ | 104 |

**Table S2.**
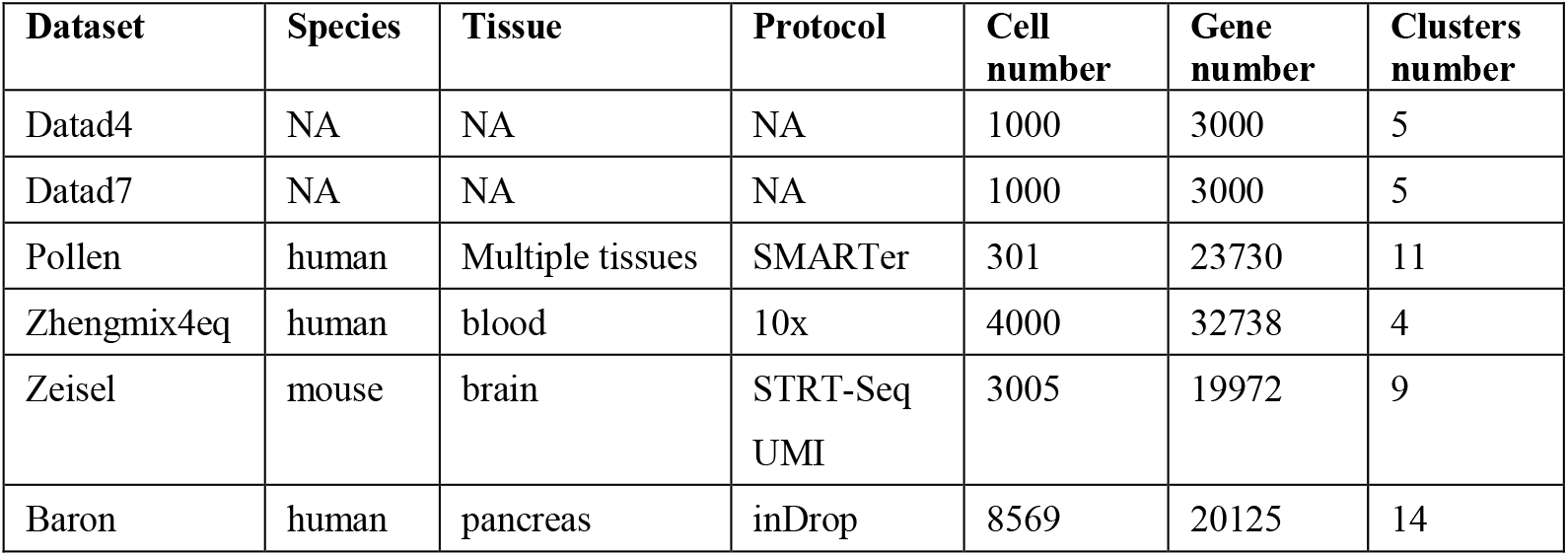
The benchmark datasets

| **Dataset** | **Species** | **Tissue** | **Protocol** | **Cell number** | **Gene number** | **Clusters number** |
| --- | --- | --- | --- | --- | --- | --- |
| Datad4 | NA | NA | NA | 1000 | 3000 | 5 |
| Datad7 | NA | NA | NA | 1000 | 3000 | 5 |
| Pollen | human | Multiple tissues | SMARTer | 301 | 23730 | 11 |
| Zhengmix4eq | human | blood | 10x | 4000 | 32738 | 4 |
| Zeisel | mouse | brain | STRT-Seq<br>UMI | 3005 | 19972 | 9 |
| Baron | human | pancreas | inDrop | 8569 | 20125 | 14 |

**Fig. S1.**
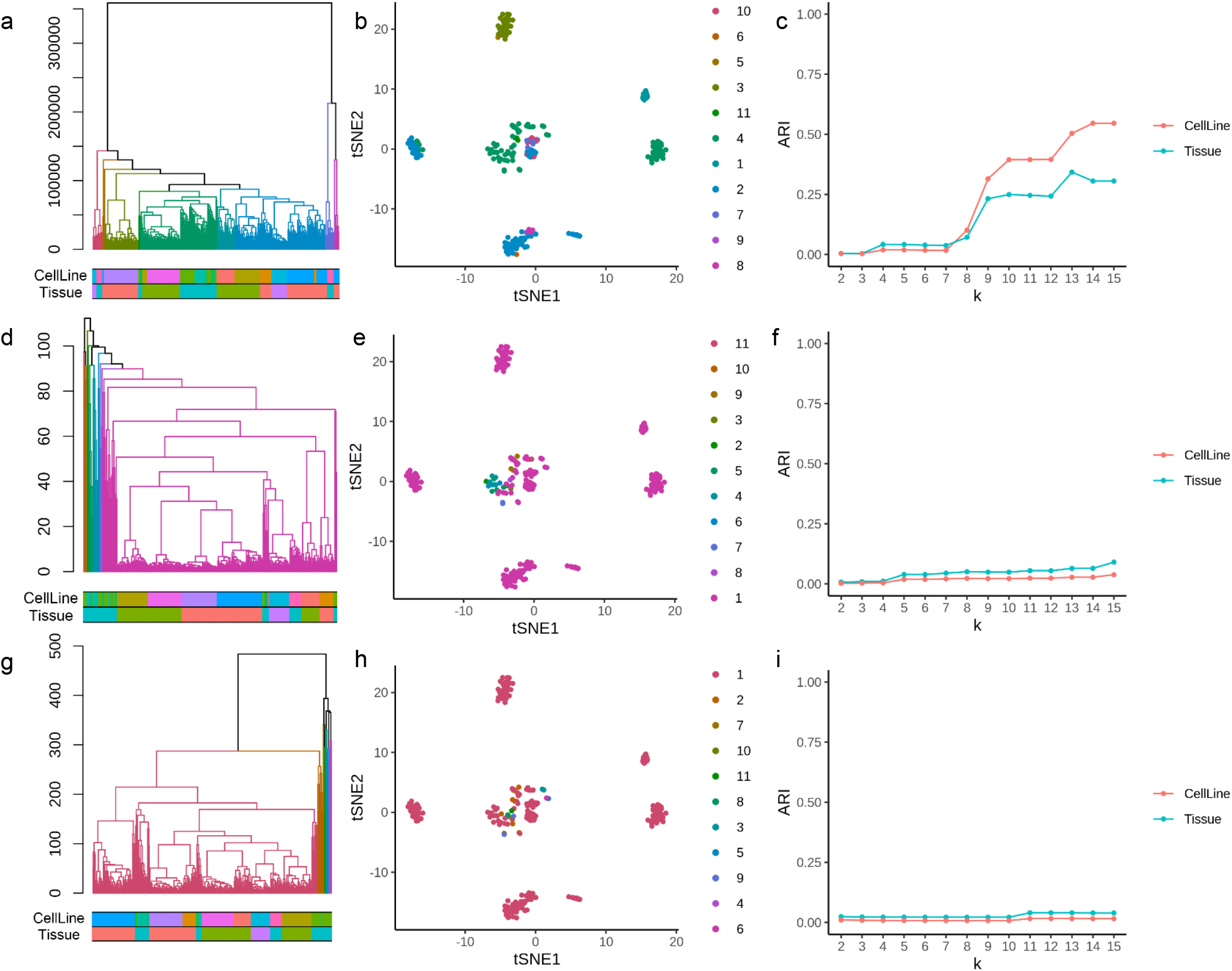
The performance of three HC-based methods in the Pollen dataset. The three rows show the results of HC, PCA+HC and GLMPCA+HC, respectively. The first column is the dendrogram. The color bars show the given labels at the cell line level and the tissue level. The second column is the tSNE plot showing the clustering result when cutting the dendrogram into 11 clusters. The third column shows the ARIs of the clustering results compared with the two labels. The x-axis is the number of clusters. It is clear that HC-based methods didn’t produce good results in Pollen dataset.

**Fig. S2.**
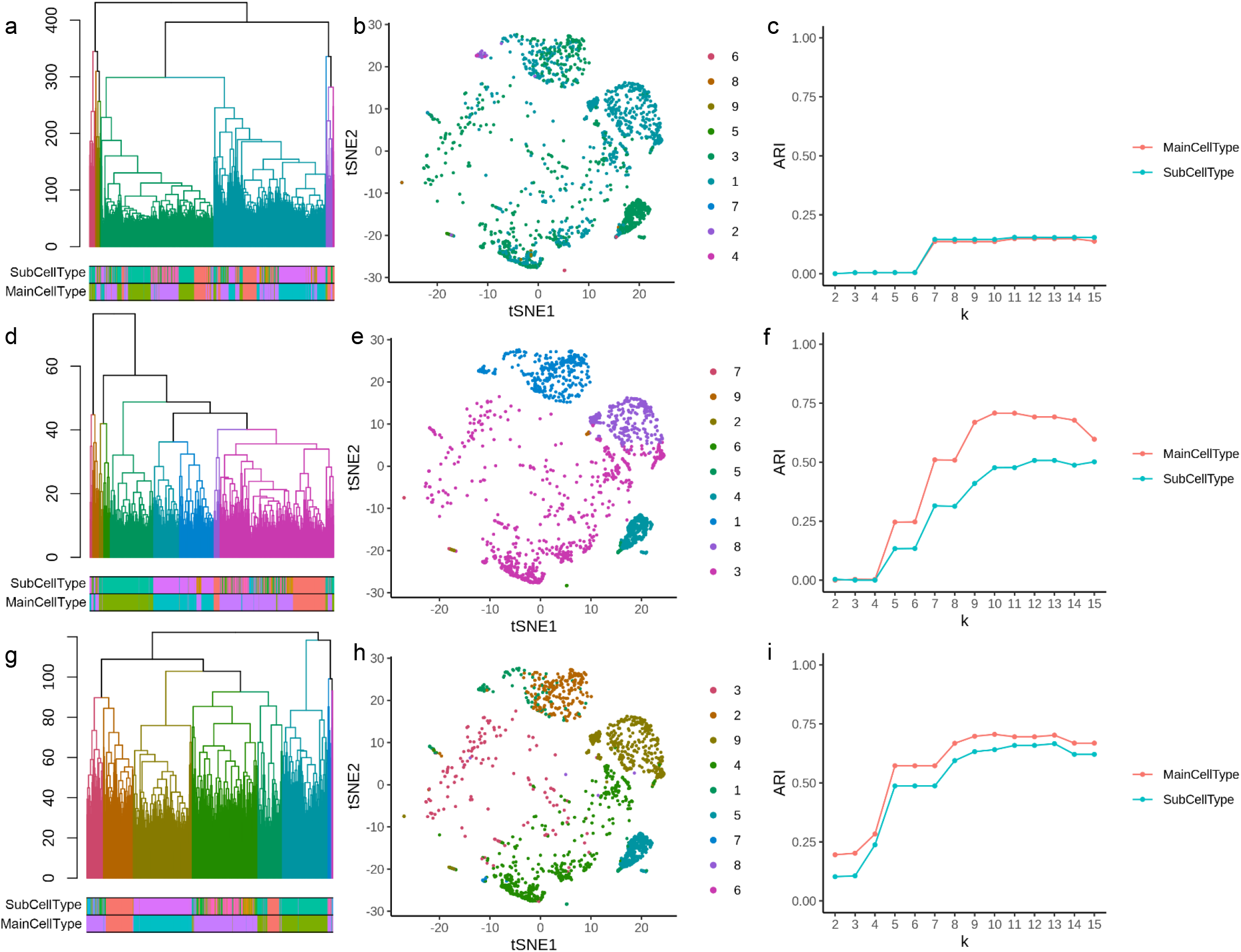
The performance of HC-based methods in the PBMC dataset. The three rows show the results of HC, PCA+HC and GLMPCA+HC, respectively. The first column is the dendrogram. The color bars show the given labels at cell line level and tissue level. The second column is the tSNE plot showing the clustering result when cutting the dendrogram into 9 clusters. The third column shows the ARIs of the clustering results compared with two labels. The x-axis is the number of clusters.

